# Heat stress reveals a fertility debt owing to postcopulatory sexual selection

**DOI:** 10.1101/2022.10.31.514482

**Authors:** Julian Baur, Martyna Zwoinska, Mareike Koppik, Rhonda R. Snook, David Berger

**Affiliations:** Department of Ecology and Genetics, Division of Animal Ecology, Evolutionary Biology Centre, Uppsala University, Uppsala, Sweden; Department of Zoology, Stockholm University, Stockholm, Sweden; Department of Zoology, Animal Ecology, Martin-Luther University Halle-Wittenberg, Halle (Saale), Germany

**Keywords:** Temperature, postcopulatory sexual selection, sperm competition, fertility, heat shock, mating system, sex, climate warming, environmental change

## Abstract

Climates are changing rapidly, demanding equally rapid adaptation of natural populations. Whether sexual selection can aid such adaptation is under debate; while sexual selection should promote adaptation when individuals with high mating success are also best adapted to their local surroundings, the expression of sexually selected traits can incur costs. Here we asked what the demographic consequences of such costs may be once climates change to become harsher and the strength of natural selection increases. We investigated how an evolutionary history of strong postcopulatory sexual selection (sperm competition) affects male fertility under acute adult heat stress. Harnessing the empirical potential of long-term experimental evolution in the seed beetle *Callosobruchus maculatus*, we assessed the thermal sensitivity of fertility (TSF) in replicated lines maintained for 68 generations under three alternative mating regimes manipulating the opportunity for sexual and natural selection. We find that males evolving under strong sexual selection suffer from increased TSF, and that male success in sperm competition (P2: sperm offense) is genetically correlated to increased TSF. Interestingly, females from the regime under strong sexual selection, who experienced relaxed selection on their own reproductive effort, had high fertility in benign settings but suffered increased TSF, like their brothers. This implies that female fertility and TSF evolved through genetic correlation with reproductive traits sexually selected in males. Paternal but not maternal heat stress reduced offspring fertility with no evidence for adaptive transgenerational plasticity among heat-exposed offspring, indicating that the observed effects may compound over generations. Our results suggest that trade-offs between fertility and traits increasing success in postcopulatory sexual selection can be revealed in harsh environments. This can put polyandrous species under increased risk during extreme heat waves expected under future climate change.

**IMPACT STATEMENT:** How will populations respond to a warming world? Of increasing concern are negative effects of elevated temperatures on fertility, which in many species are observed for temperatures substantially lower than the ones causing death. Incorporating knowledge on species-specific thermal fertility limits has improved estimates of current species’ ranges but renders a more pessimistic view of the potential for adaptive responses under climate change. Sexual selection is a process that can interact with thermal sensitivity of fertility and is strongest in males of polyandrous species, in which females mate multiply and sperm of multiple males compete for fertilization of female eggs. Therefore, males of polyandrous species often invest heavily in sperm competition. However, given finite resources, increased investment in sperm competition can come at an expense of other processes needed to maintain the integrity of the male germline, which when compromised can reduce fertility and offspring quality. How may such male investment, fuelled by sexual selection, affect species responses to climate warming? To address this question, we first evolved populations under different laboratory settings that independently manipulated the levels of natural and sexual selection. We exposed adults from these populations to acute heat stress and measured the fertility of males and females. We find that sexual selection on males leads to a fertility debt that is revealed under heat stress. This debt was also apparent in females, who themselves were not selected for increased reproductive investment. Thus, genes under sexual selection in males seems to have impaired fertility in both sexes under heat stress. Forecasts of species response to climate change that do not incorporate thermal fertility limits and sexual selection may therefore underestimate species vulnerability to increasing temperatures.

## INTRODUCTION

Sexual selection is a strong evolutionary force that can select for traits that are associated with considerable costs in the face of natural selection (Andersson, 1994; Zahavi, 1975). Harsh environments that impose strong natural selection are therefore predicted to limit the evolution of excessive expression of sexually selected traits in favour of allocation to maintenance and survival (Buchanan, 2000; Candolin & Heuschele, 2008; Zahavi, 1975). It is well recognized that anthropogenic environmental change is placing many organisms under the threat of extinction by imposing severe challenges on natural populations (IPCC 2022). How may such rapid increases in the force of natural selection affect species that have had a long-term history of evolving under strong sexual selection? Intuitively, one might expect polygamous species that invest heavily in sexually selected traits to suffer severe fitness losses when environments change to become harsher and impose greater needs for allocation to maintenance. Moreover, because sexually selected traits are often linked to fertility, these consequences could be severe, as fertility is typically highly sensitive to environmental stress and a strong determinate of population-level viability (Parratt et al., 2021; Walsh et al., 2019).

In polyandrous species, postcopulatory sexual selection through sperm competition represents a central part of the selective process (Birkhead & Pizzari, 2002). Sperm competition can lead to the evolution of increased sperm numbers (Boschetto et al., 2011; Simmons & Fitzpatrick, 2012; Wedell et al., 2002) and investment into, presumably costly, sperm traits such as swimming velocity (Gage et al., 2004), flagellum length (Godwin et al., 2017), and ornamentation (Lüpold et al., 2016; Silva et al., 2019). However, the need for more numerous and competitive sperm also requires increased maintenance to sustain the integrity of the germline and ensure high fertility and offspring quality (Dowling & Simmons, 2009; Monaghan & Metcalfe, 2019). Such maintenance, including DNA repair, antioxidant defence and apoptosis, is tied to considerable costs (Chen et al., 2020; Kirkwood, 2005; Kirkwood et al., 1979; Lemaître et al., 2020; Maklakov & Immler, 2016). Hence, if organisms balance investment into sperm competition against germline maintenance, increased demand on maintenance under rapid environmental change could cause a severe reduction of male fertility in species with intense postcopulatory sexual selection.

Climate warming and the incidence of heat waves is one of the most common and impactful consequences of anthropogenic environmental change (Bathiany et al., 2018; IPCC, 2022; Johnson et al., 2018; Varela et al., 2020), and male fertility is highly sensitive to increased temperatures (Chirgwin et al., 2021, 2020; Iossa, 2019; Rodrigues et al., 2022; Sales et al., 2021; Vasudeva et al., 2019; Walsh et al., 2019; Wang and Gunderson, 2022). We therefore tested the general prediction that a history of strong sexual selection might lead to greater environmental sensitivity of male fertility by investigating how experimental evolution under different levels of natural and sexual selection affects the thermal sensitivity of fertility in the seed beetle, *Callosobruchus maculatus*, a model species for studies on postcopulatory sexual selection. We used a set of replicated lines that had evolved for 68 generations under three alternative mating regimes, manipulating the relative strengths of sexual and natural selection (natural selection only (N), natural and sexual selection (N+S), or sexual selection only (S)) in benign and constant laboratory conditions. Previous work has shown that these regimes have evolved differences in a variety of reproductive phenotypes. For example, populations evolving under strong sexual selection and minimized natural selection show greater postcopulatory reproductive success (Koppik et al., 2022) and different sperm allocation patterns with more plastic germline maintenance (Baur & Berger, 2020). Hence, we predicted that these males would suffer increased thermal sensitivity of fertility (henceforth: TSF) compared to males that have evolved without sexual selection.

Plastic male allocation decisions in response to social cues, such as the presence of receptive females or male competitors, has been observed in several polyandrous taxa (e.g., Bretman et al., 2011, 2010; Ramm and Stockley, 2009). If such plasticity shifts resources away from germline maintenance in favour of investment in sperm competition, this could likewise reduce TSF under the trade-off scenario. Indeed, the studied evolution regimes of *C. maculatus* have previously been shown to differ in how socio-sexual interactions affect plastic changes in ejaculate traits and germline maintenance (Baur & Berger, 2020; Koppik et al., 2022). We therefore also explored the direct effects of the presence of male rivals on the plasticity of TSF across the three evolution regimes, predicting that such interactions would generally reduce male TSF.

Female fertility is typically a stronger limiting factor on population growth rate than male fertility (Caswell, 2006; Manning, 1984). Understanding if and how female fertility is moulded by the mating system is therefore important for predicting demographic consequences under future climate warming (e.g., Fox et al., 2019). Females can exhibit complex trade-offs between reproduction and maintenance (Harshman & Zera, 2007) and often evolve costly counter-adaptations to male mating strategies (Andersson & Simmons, 2006; Arnqvist & Rowe, 2002; Pizzari & Snook, 2003; Rönn et al., 2007; Ryan, 1998) and presumably costly mechanisms that allow them to exert cryptic female choice of male sperm (W. Eberhard, 1996; W. G. Eberhard & Cordero, 1995; Shuker & Simmons, 2014; Telford & Jennions, 1998). However, it remains unclear how such female adaptation to mating interactions affect their stress tolerance. Additionally, because male and female reproductive traits often share a genetic basis, it is possible that male adaptation could target genes that also affect female reproduction and maintenance. To explore how sexual selection affects female fertility responses to environmental stress we therefore also assayed TSF in females from the three evolution regimes.

The severity of the impact of environmental stress on population viability depends on if and how effects on fertility are carried over to subsequent generations. However, it remains unclear whether such transgenerational effects typically confer adaptive or detrimental responses in offspring (Bonduriansky & Day, 2009; Donelson et al., 2018; A. A. Hoffmann & Sgrò, 2011). Indeed, evidence for whether parental heat stress positively or negatively affects offspring TSF is scarce but seemingly indicates that heat stress experienced by parents reduces offspring TSF (Burgess & Marshall, 2011; Diaz et al., 2021; Uller et al., 2013). Population-level consequences of reductions in fertility should also depend on which sex that is most severely impacted (Caswell, 2006; Manning, 1984), but studies assessing the sex-specificity of transgenerational effects of heat stress are rare. We therefore also assessed sex-specific transgenerational effects on TSF.

Our results support the hypothesis that previous male adaptation under strong directional sexual selection on sperm traits may lead to detrimental effects on fertility once temperatures rise, and that these effects may permeate through generations. We also find evidence suggesting that sexual selection may lead to increased thermal sensitivity of female fertility, most likely via genetically correlated responses to selection on males. Forecasts of responses to environmental change should thus incorporate sexual selection and the mating system to accurately predict species vulnerability.

## METHODS

### Predictions for how a history of strong sexual selection affects the environmental sensitivity of male fertility

To formalize general predictions for how an evolutionary history of strong postcopulatory sexual selection affects the environmental sensitivity of male fertility, we employed a life history theory framework and the “Y-model” for allocation and acquisition trade-offs (de Jong & van Noordwijk, 1992; Houle, 1991). The model is described in full in Supplement S1. In brief, the model assumed that “fitness” is the product of competitive fertilization success and gamete viability. It was further assumed that individual condition (*C*) determines the amount of resources that can be allocated to germline maintenance (*M*) in form of anti-oxidative defence and repair needed to maintain ejaculate quality and gamete viability (Dowling & Simmons, 2009; Friedberg et al., 2005) and reproductive effort (*R*) in form of gamete production and ejaculatory components that increase a male’s competitive fertilization success, such that: *C = R + M*. This results in an allocation trade-off between reproductive effort and germline maintenance that impacts sperm competition success and fertility, respectively.

Competitive fertilization success was modelled as an increasing power function of reproductive investment (*R*), with the strength of postcopulatory sexual selection given by exponent *b*, with higher values of b indicating stronger sexual selection. Gamete viability was modelled as an increasing power function of allocation to maintenance (*M*), with the strength of viability selection in a given environment given by exponent *a*, where higher values of *a* indicate harsher conditions. Optimal allocation between reproductive effort and maintenance was found by maximizing fitness for different values of *a* and *b* (Fig. 1A). We then calculated the fertility reduction resulting from abrupt increase in environmental harshness (increase in *a*) for species with alternative allocation strategies corresponding to differences in the relative strengths of sexual selection (different values of *b*) and viability selection (different values of *a*) in their ancestral environment (Fig. 1B).

**Figure 1:**
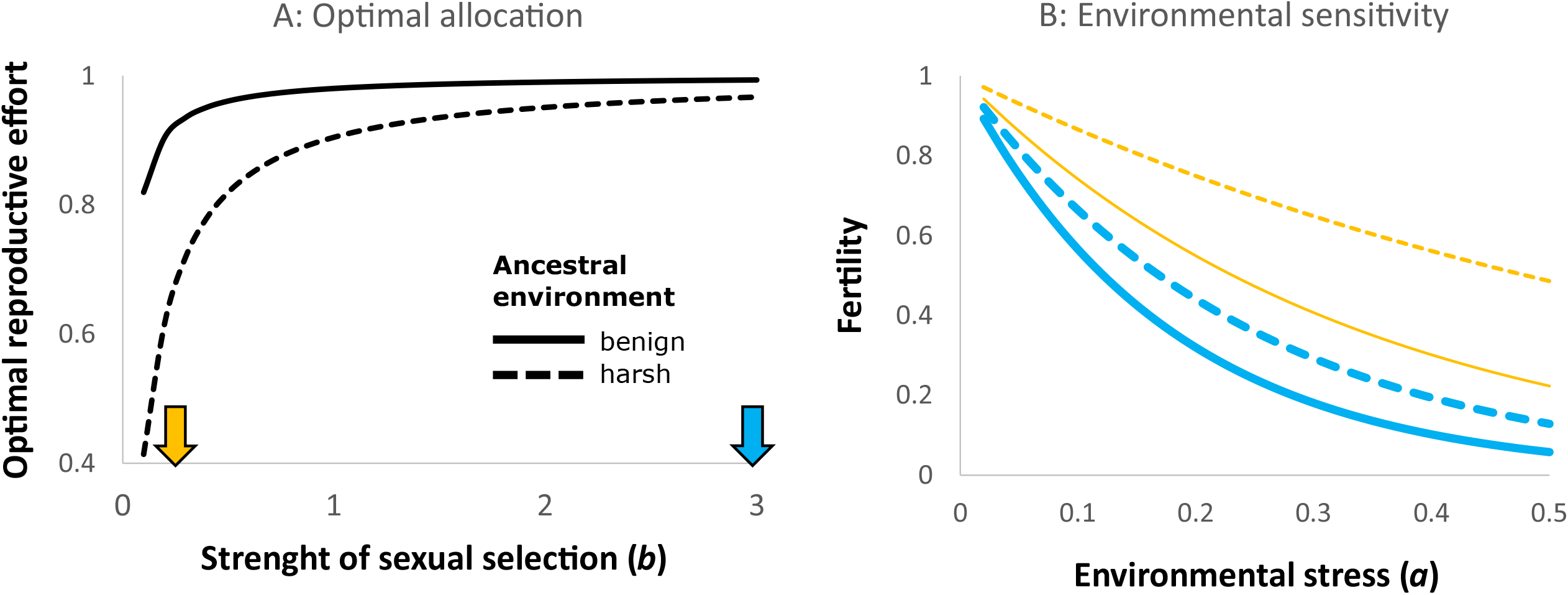
The expected relationship between the strength of postcopulatory sexual selection and fertility under environmental stress. In A) the optimal fraction of resources devoted to reproductive effort (traits that increase sperm competition success) is shown for different strengths of postcopulatory sexual selection in an environment that is either benign and imposes weak viability selection (*a* = 0.02, full line) or relatively harsh (*a* = 0.10, hatched line). The downwards facing arrows along the x-axis depict hypothetical species that have evolved to maximize fitness under either weak (*b* = 0.2, yellow) or strong (*b* = 3.0, blue) postcopulatory sexual selection. In B) the consequences for fertility are given for the different scenarios once an abrupt environmental change occurs that increases viability selection (via exponent *a*). Postcopulatory sexual selection promotes investment into reproductive effort, which leads to strong declines in fertility once the environment becomes stressful. Polyandrous species evolving in benign ancestral conditions, that devote most resources to reproductive effort and least to maintenance, are predicted to be most at risk (full blue line).

### Study species

*C. maculatus* originates from the tropical and subtropical regions of the world and is a common pest of stored fabaceous seeds. Females glue their eggs on host beans and the larvae develop inside the beans for roughly three weeks before eclosing as sexually mature adults (C. W. Fox, 1993). Reproduction starts a few hours after eclosion and usually takes place within the first few days of adulthood (C. W. Fox, 1993). The adult life span of *C. maculatus* typically ranges between 7 and 12 days under aphagous conditions, with females living longer than males. *C. maculatus* is frequently used as a model system to study sexual selection and sexual conflict (Arnqvist et al., 2021; Baur et al., 2019; Berger, You, et al., 2016; Bilde et al., 2009; Dougherty et al., 2017; Eady, 1995; Gay et al., 2009; Lieshout et al., 2013; Rönn et al., 2006, 2007) because males are known to compete fiercely over access to females, leading to high levels of promiscuity and postcopulatory sexual selection. Sperm regeneration rates have been shown to evolve in response to sexual selection in *C. maculatus* (Baur & Berger 2020) and to be associated with increased metabolic expenditure (Immonen et al., 2016). Female beetles show a noticeable kicking behaviour upon a mating attempt by a male, and potentially cryptic female choice (Lieshout et al., 2014). While preferred temperatures range from 25°C to 30°C (C. W. Fox et al., 2006; Martinossi-Allibert et al., 2017; Vasudeva et al., 2014), several experiments indicate that this species exhibits tolerance to even higher temperatures (Berger et al., 2017, 2021; Lale & Vidal, 2003; Loganathan et al., 2011).

### Experimental evolution regimes

The experimental evolution lines were created from a stock population sampled from a natural population in Lomé, Togo (06°10#N 01°13#E), in 2010. Previous studies have demonstrated that this genetic stock harbours substantial standing genetic variation for behaviour, life history and sex-specific reproductive success (Berger, Martinossi-Allibert, et al., 2016; Berger, You, et al., 2016; Grieshop et al., 2021; Grieshop & Arnqvist, 2018). The three experimental evolution regimes (outlined below) have been studied extensively and show divergence in a range of reproductive traits, including sex-specific competitive reproductive success (Martinossi-Allibert, Thilliez, et al., 2019), mating behaviour (Baur et al. 2019), germline maintenance (Baur and Berger, 2020; Koppik et al. 2022), postcopulatory reproductive success (Koppik et al., 2022) and immunity (Bagchi et al., 2021). Three replicate lines were started per evolution regime, but one line was lost for the S regime prior to experiments. Each line was maintained at an effective population size of approximately 150. For more details on the selection protocol see (Martinossi-Allibert, Thilliez, et al., 2019).

#### N + S (natural and sexual selection)

This regime was designed to resemble the natural mating system of *C. maculatus*. This regime allows for pre- and postcopulatory sexual selection as well as viability and fecundity selection as males and females were freely interacting during the entire reproductive and egg-laying period.

#### N (only natural selection)

Under this regime, a virgin male and female were paired at random to form monogamous couples, removing sexual selection while allowing for viability and fecundity selection. This regime should select for males with ejaculatory components with beneficial effects on female fertility.

#### S (only sexual selection)

This mating regime was designed to allow for sexual selection on males while attempting to minimize natural selection, thereby removing genetic constraints on the evolution of secondary sexual characters (i.e., sperm traits) imposed by natural selection on correlated traits expressed in both sexes. Males and females were first allowed to mate and interact freely (i.e., sexual selection proceeded) for 48h without egg-laying substrate for females, after which all females were collected in individual 60 mm petri dishes, each containing roughly 30 beans onto which the females could oviposit. Exactly one male and one female beetle per dish were picked to contribute to the next generation. This effectively removed selection on female fecundity by making sure that each female contributed equally, and only two offspring, to the next generation (offspring numbers per female typically range between 50-100). We note that although viability selection was not actively prevented in this regime, female mortality was very low (1 out of 100 females died every 1-2 generations) and egg-to-adult survival is high (>95%) in all lines, thus, viability selection is unlikely to be effective in this or any other regime.

### Assessing male and female TSF via heat shock

Following 68 generations of experimental evolution, all lines were maintained for two generations under common garden conditions (typical laboratory conditions resembling the N+S regime; See Supplement S2 for a graphical illustration of the experimental design). The experiment was performed in two blocks, each consisting of three experimental days coinciding with the peak emergence of the beetles in each block. We picked virgin focal adults from all lines within 24 hr after eclosion. Virgin focal males and females were isolated in perforated 0.5 ml Eppendorf tubes for 24 hr (isolated treatment), except the males assigned to compete (male-male treatment). These males were placed in 35 mm petri dishes in groups of three. After 24 hrs, we randomly selected half of the beetles in each group for heat shock exposure. Prior to the heat shock, we moved all males from the male-male treatment individually into perforated 0.5 ml Eppendorf tubes, to ensure the same conditions during the heat shock. The heat shock consisted of 20 minutes in an incubator at 55°C at high relative humidity, which has previously been shown to result in a reduction of fertility while remaining in a range that is ecologically relevant for *C. maculatus* (Baur et al., 2022). To confirm the ecological relevance of the selected temperature, we used NicheMapR (Kearney & Porter, 2017) to run a microclimate model assuming a global temperature increase of 1.5*°*C. The model showed that soil temperature can reach up to 70*°*C and air temperatures up to 50*°*C in Lomé, Togo, placing 55*°*C in a range that a ground-dwelling insect may experience (see Supplement S2 for the NicheMapR model). We mated all focal beetles to reference individuals of the opposite sex (untreated beetles from the ancestral stock population) starting 20 minutes after the heat shock. We mated heat shocked males from the isolated treatment a second time, seven hours after heat shock, to investigate time effects on TSF. All matings were performed in 60 mm petri dishes on a heating plate at benign 29°C. We excluded 37 couples that did not mate within 75 minutes. This resulted in a total of 1123 mating couples for which fertility (number of adult offspring) was recorded, with on average 47 couples per experimental cell (a combination of: sex, heat shock treatment, and male social treatment; for exact sample sizes see Supplement S4).

We did not mate untreated control males a second time in our experiment, since males of *C. maculatus* are able to mate multiply without detectable declines in fertility (Rönn et al., 2008). Hence, there was no formal control group for heat shocked males that mated a second time, 7h after the heat shock application. We therefore confirmed that our results (see below) were indeed caused by responses to heat stress, and not an effect of mating order or male aging *per se*, by performing a follow-up experiment comparing fertility from first and second matings of untreated male beetles from the S regime. More detailed methods and the results from this experiment are summarized in Supplement S5.

### Transgenerational effects

We investigated transgenerational effects using the three lines from the N+S regime, which is closest to this species’ natural polygamous mating system. We studied effects on F1 offspring fertility and TSF (son, daughter, or neither one, was heat shocked) from heat shock applied to the F0 parents (mother, father, or neither one, was heat shocked) in a fully crossed design (see Supplement S6 for a graphical illustration of the design). F0 males belonged to the isolated treatment in the original experiment, and all F1 offspring derived from the first mating following heat shock in the F0. All focal F1 offspring were mated to untreated partners originating from the parental control treatment and same experimental evolution line.

### Statistical analyses

We used Bayesian generalized mixed effects models implemented in the package MCMCglmm (Hadfield, 2010) for R (R Core Team, 2020). Ggplot2 was used for graphical illustration (Wickham, 2016).

We performed three main analyses including the experimental evolution lines. We used uninformative and weak priors and included line identity crossed with the applied treatments as random effect terms. The 6 experimental days (2 blocks with 3 days each), crossed with the heat shock treatment, were included as additional random effect terms. First, we tested for effects of evolution regime on TSF from the first and second mating in isolated males in a model that incorporated fixed effects of the heat shock treatment, mating number, and evolution regime, as well as all higher order interactions (see Supplement S7). This model was including only data on isolated males (first and second ejaculate). Second, we tested for effects of the presence of male rivals on the TSF from the first mating (males from the male-male treatment were not assayed for the second mating) in a model including the fixed effects of heat shock treatment, evolution regime, male social treatment, as well as all higher-order interactions (see Supplement S8). Third, we analysed effects of evolution regime on female TSF in a model that included heat shock treatment and evolution regime (all females were kept isolated prior to heat shock and were all allowed to mate only once following it, see Supplement S9 for model details)). When analysing effects of heat shock in males in the first model including both the first and second mating, we excluded four males that failed to mate during the first census time. All these males derived from the third day of the second block and were limited to the heat shock treatment. When analysing the effects of male rivals on fertility in the second model, we excluded these four isolated males and an additional six males from the male-male treatment that also failed to mate after the heat shock. The number of excluded males was small and equally distributed over the evolution regimes, and their inclusion/exclusion did not affect results qualitatively (for an analysis including males that did not mate, with their fertility of zero, see Supplement S8 and S10). The model used to investigate transgenerational effects included the sex of the focal individual from the parental generation, the heat shock treatment of the focal parental individual (heat shock, control), and the treatment of the F1 couple (male heat shock, female heat shock, control) as interacting fixed effects. Line identity was also included as fixed effect. The experimental block from the parental generation and its interaction with the sex and treatment of the focal parental individual were included as random effects.

We assumed a Poisson distributed error for the response in all models used for testing of statistical significance. For illustrative purposes, we also calculated the effect size as: TSF = 1 – mean number of offspring_heat shocked_ / mean number of offspring_control_ (thus giving the proportional reduction in offspring produced attributed to heat shock), based on posteriors from models equivalent those described above, but using a Gaussian response. The results of these Gaussian models are presented in Figures 2-5 and for all offspring numbers or fertility reductions (incl. TSF) reported in the result section. The mean number of offspring was high (range 55-85 for the different treatments), so the response was approximately normal, and the resulting estimates from the Poisson and Gaussian models were qualitatively identical. We ran our models for 2.2M iterations with an initial burn in of 200k iterations and a thinning factor of 2000 to avoid autocorrelations, resulting in 1000 uncorrelated posterior samples from which posterior means, 95% credible intervals and two-tailed P-values were calculated based on model posterior distributions (model specifications in Supplements S7-9, S11).

**Figure 2:**
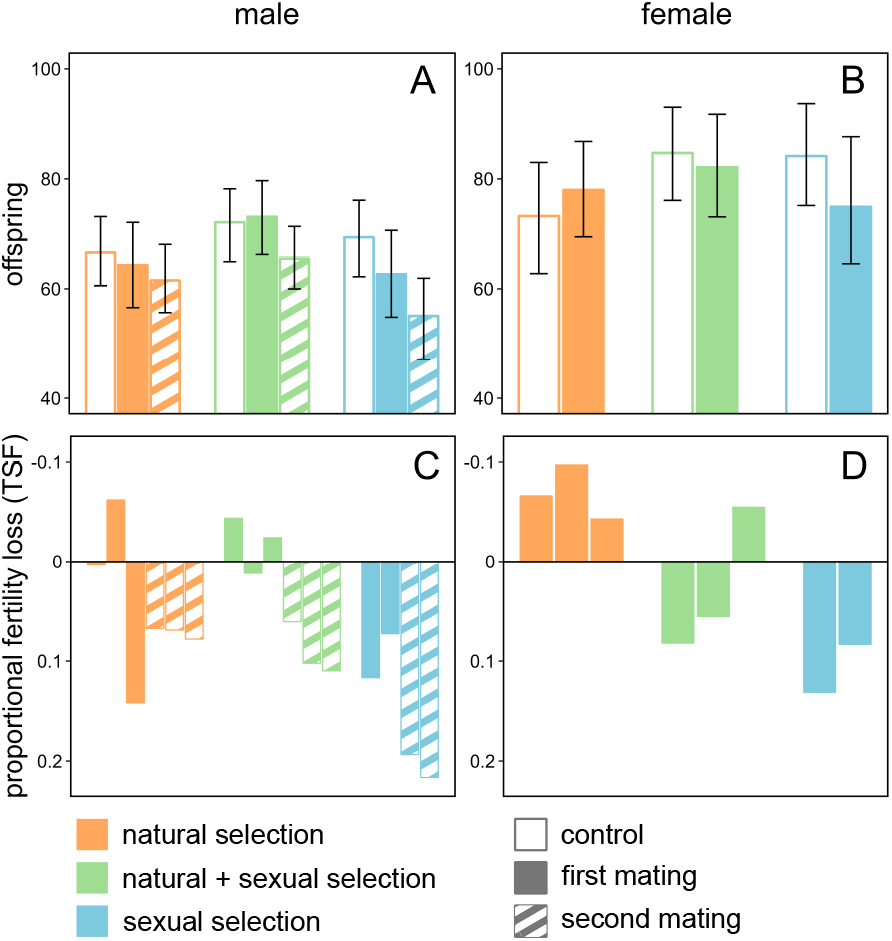
The evolution of male and female TSF under varying levels of natural and sexual selection. The top panels show fertility of focal (isolated) males (A) and females (B) that were either heat shocked (closed bars, first ejaculate/mating 20 minutes after heat shock; striped bars, second ejaculate 7 jours after heat shock) or controls (open bars, first ejaculate/mating), originating from the N (natural selection only: orange), N+S (natural and sexual selection: green), or S (sexual selection only: blue) regime. Bars represent posterior means and whiskers 95% credible intervals. The bottom panels show TSF for experimental evolution regimes, with male TSF (C) and female TSF (D) shown separately for each replicate line. TSF was calculated as 1-(offspring_heat shocked_/offspring_control_), using raw data means. Hence, positive values indicate reductions in offspring due to heat shock.

To explore possible mechanistic links between the evolution of sexually selected postcopulatory reproductive traits and male TSF, we made use of previously published data from the same eight replicate lines as used here on sperm production (males were mated three times within 90 minutes to deplete sperm storage, 25 hours later males were allowed to mate again, sperm production refers to the increase in sperm number per ejaculate between the third and fourth mating), following 29 generations of experimental evolution (Extended data Fig. 5 in Baur & Berger 2020), and postcopulatory reproductive success in form of sperm defence (P1, i.e., the focal male is the first of two males to mate with the female) and sperm offense (P2, i.e., the focal male is second of two males to mate with the female), following 51 generations of experimental evolution (Fig. 1 in Koppik et al. 2022). We then estimated genetic correlations between these traits and the fertility reduction induced by heat shock (i.e., the TSF) in the second mating, based on line means. P1 and P2 were logit-transformed before analysis, as these traits are proportions ranging between 0 and 1.

## Results

### Predictions for how a history of strong sexual selection affects the environmental sensitivity of male fertility

Increased allocation to sperm competition traits at the expense of germline maintenance is predicted to evolve when the expected fitness return of increased postcopulatory reproductive effort (given by exponent *b*) is high but is disfavoured in harsh environments that impose strong viability selection on gametes (given by exponent *a*) (Fig. 1A). Once environmental harshness suddenly increases (*a* increases) fertility declines, but more so in species that have evolved their optimal allocation strategy in benign environments (small ancestral *a*). For any strength of viability selection (*a*) in the ancestral environment, populations that have evolved under a history of strong sexual selection (high *b*) are predicted to suffer a greater fertility loss following increased environmental stress (Fig. 1B). We note that this simple model does not consider several conditions that could alter response, such as: adaptive germline plasticity, further evolution in response to the change in environmental harshness, or how changes in environmental harshness may cause changes in the strength of postcopulatory sexual selection (*b*) itself (e.g., Martinossi-Allibert et al., 2019a; Svensson and Connallon, 2019).

### Effects of evolution under different mating regimes on the thermal sensitivity of male fertility

To test this general prediction, we compared the TSF of isolated males from the three evolution regimes that differed in the strength of postcopulatory sexual selection. For untreated (control) males, we found no fertility differences between the evolution regimes, and thus no evidence for a general reduction in the fertility of S males that had evolved under sexual selection in the absence of natural selection (mean number of offspring produced with 95% CI: N: 66.4 [58.8, 73.4]; N+S: 71.8 [64.8, 78.8]; S: 69.0 [62.9, 77.7]; all pairwise p_MCMC_ > 0.16, Fig. 2A). For naïve males kept in isolation prior to the heat shock treatment, heat stress had significant effects on fertility in the second, but not in the first mating (first mating: p_MCMC_ = 0.116, second mating: p_MCMC_ = 0.028). Furthermore, in accordance with predictions (Fig. 1B), the reduction in fertility caused by heat shock was strongest in the S regime (TSF first mating: N: 0.03 [-0.12, 0.16]; N+S: -0.01 [-0.14, 0.10]; S: 0.09 [-0.04, 0.22]; TSF second mating: N: 0.07 [-0.05, 0.20]; N+S: 0.09 [-0.02, 0.21]; S: 0.21 [0.08, 0.33], regime:treatment interaction for second mating; N vs. N+S: p_MCMC_ = 0.76, N vs. S: p_MCMC_ = 0.026; N+S vs. S: p_MCMC_ = 0.014, Fig 2A, C) (Supplement S7).

### Effects of male rivals

We explored whether potential plasticity in germline allocation in response to cues from male rivals affected male TSF. Male-male interactions resulted in overall negative effects on fertility in untreated (control) males (fertility reduction, mean offspring produced with 95% CI: 5.6 [2.3, 8.7], p_MCMC_ = 0.015; Fig. 3; see Supplements S8 and S10 for the analysis including males that failed to mate), suggesting that these interactions had costs. However, when analysing the regimes separately, we found an effect of male-male interactions only in N+S control-males (fertility reduction: 8.6 [2.0, 14.7], p_MCMC_ = 0.028), and S control-males (fertility reduction: 6.4 [1.6, 11.1], p_MCMC_ = 0.033), whereas there was no effect in N control-males (fertility reduction: 1.7 [-4.1, 7.0], p_MCMC_ = 0.69) (Fig. 3). This pattern suggests that N males may have evolved to invest less into male-male competition under the removal of sexual selection, although we did not find a statistically significant difference in the effect of male-male interactions between regimes (interaction: N vs N+S: p_MCMC_ = 0.083, N vs S: p_MCMC_ = 0.23). However, there was no suggestion that male-male interactions worsened the impact of heat shock on male fertility (all interactions: p_MCMC_ > 0.08, Fig. 3). We note that the effect of male-male interactions on fertility was only investigated in the first mating, where effects of heat shock were overall very modest, which may have reduced the chances of detecting such effects.

**Figure 3:**
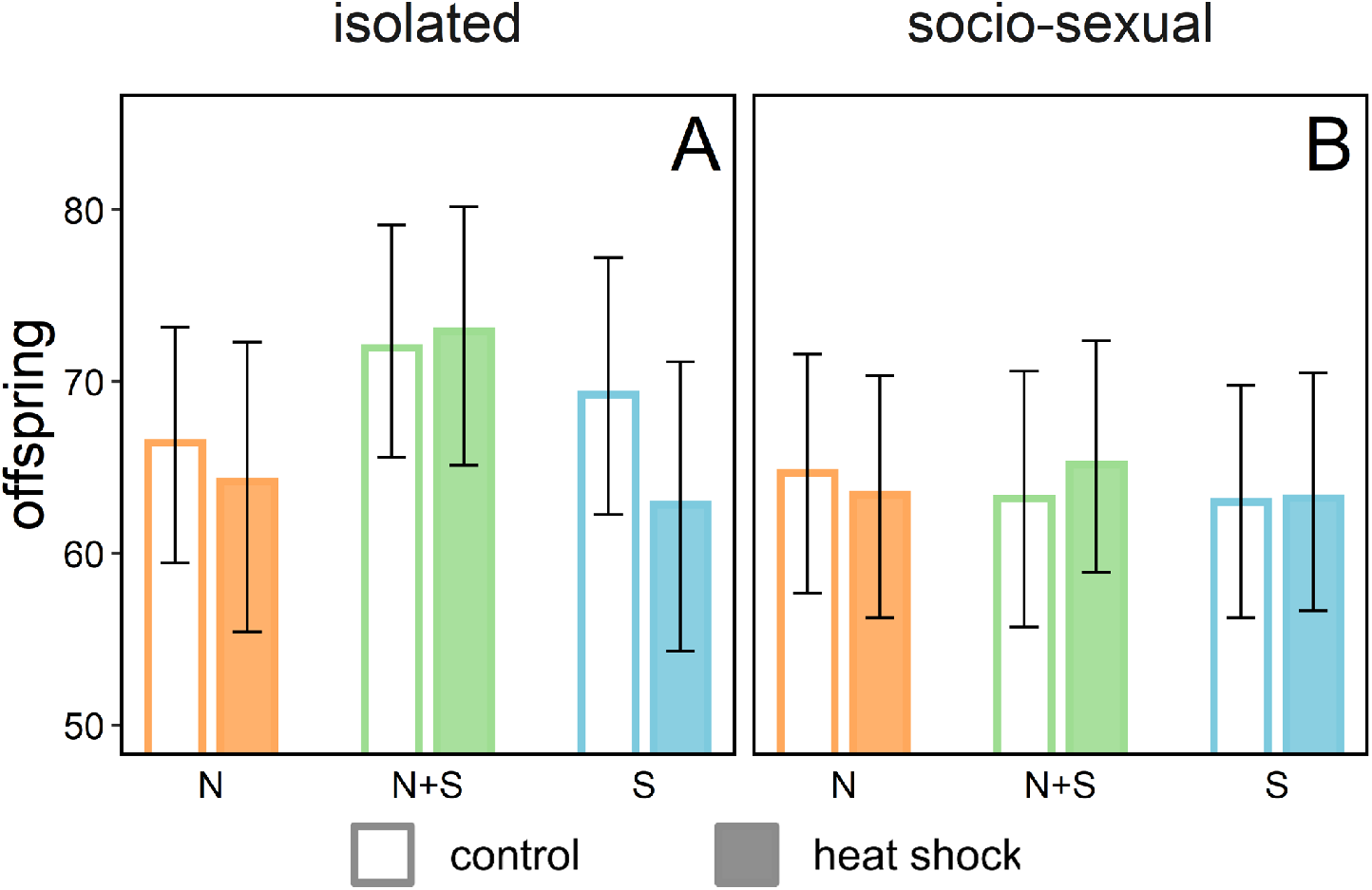
The effect of male-male interactions on male fertility. Fertility of couples from the N (natural selection only: orange), N+S (natural and sexual selection: green), or S (sexual selection only: blue) regime. Focal males were either kept at benign conditions (open bars) or exposed to heat shock (closed bars) and were either kept isolated (A) or in groups of three (B) prior to heat shock and mating. Bars represent posterior means and whiskers 95% credible intervals.

### Mechanistic links between investment in sperm competition and thermal sensitivity of male fertility

We found a strong and statistically significant correlation between male TSF (assayed in the second mating) and previously reported estimates of male success in sperm offense, P2 (r = 0.89, p = 0.003), but not for sperm defence, P1 (r = 0.39, p = 0.34) nor sperm production (r = 0.03, p = 0.95), implying that improvement in a male’s sperm offense is associated with increased sensitivity to thermal stress (Fig. 4).

**Figure 4:**
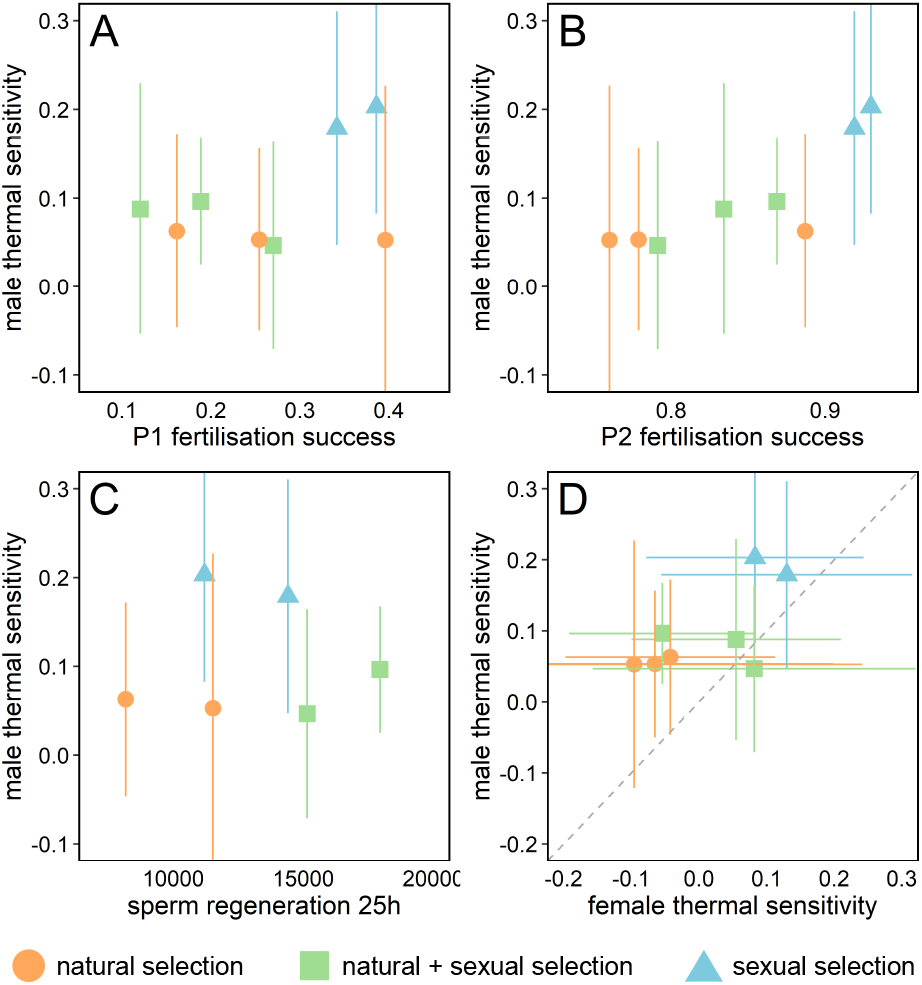
Correlations between male TSF and postcopulatory traits: (a) sperm defence (P1 fertilisation success), (b) sperm offense (P2 fertilisation success) (P1 & P2 data from Koppik et al., 2022), (c) sperm regeneration (measured as the number of sperm transferred in mating, 25 hours after sperm depletion; data from Baur & Berger, 2020), and also (d) female TSF. P1 and P2 were logit-transformed before analysis. Individual data points represent line means for N (orange circles), N + S (green squares), and S lines (blue triangles). The diagonal line in panel d represents equal male and female TSF, indicating that male TSF is higher compared to female TSF. Whiskers represent 95% Bayesian credible intervals. Note that the axes of some panels have been adjusted for better illustration, which cuts off the error bars in some instances.

**Figure 5:**
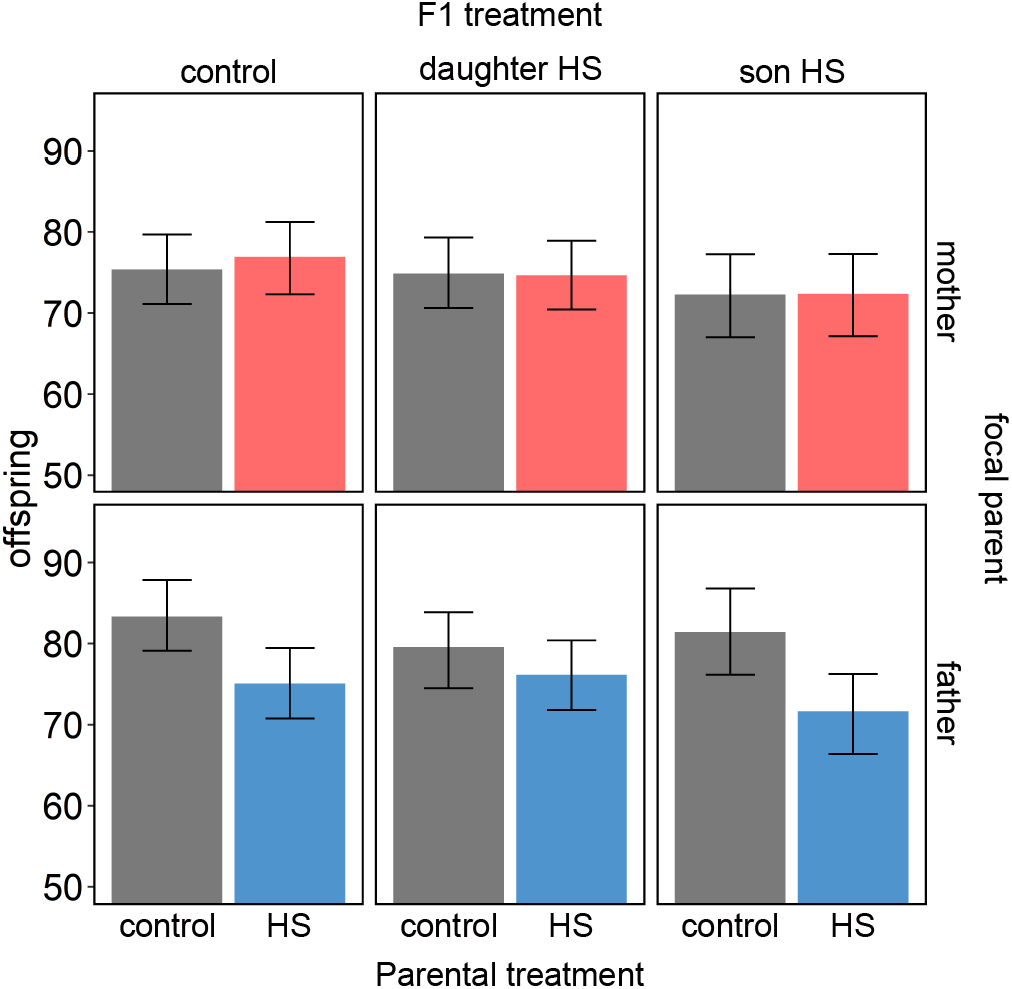
Sex-specific transgenerational effects of adult heat shock. Fertility for F1 couples in which the focal parent originates from the N+S regime (while the mating partner was a reference individual from the ancestral line)) and was either maintained at benign control temperature (grey bars) or underwent the heat shock treatment (coloured bars). F1 couples were either controls (first column) or contained a heat shocked female (second column) or a heat shocked male (third column). Mating crosses were performed while avoiding inbreeding. Bars represent posterior means and whiskers 95% credible intervals.

### Effects of evolution under different mating regimes on the thermal sensitivity of female fertility

Interestingly, the fertility of both S and N+S females, was higher than the fertility of N females in the untreated control group (mean number of offspring produced and 95% CI: N: 73.3 [62.7, 82.8]; N+S: 84.9 [76.5, 93.2]; S: 84.0 [74.8, 93.1]; N vs. N+S: p_MCMC_ = 0.056, N vs. S: p_MCMC_ = 0.032) (Fig. 2B, D). These results thus suggest that females have evolved increased reproductive output as a result of sexual selection on their brothers, demonstrating that the removal of natural (fecundity and viability) selection in the S regime has not led to any detectable decline in fertility under benign (ancestral) lab conditions.

While there was no main effect of heat shock across all regimes (Fig. 2B, D), evolution regimes varied in the extent of TSF (N: -0.07 [-0.24, 0.08]; N+S: 0.03 [-0.08, 0.15]; S: TSF: 0.10 [-0.01, 0.25], interaction regime:heat shock, N vs. S: p_MCMC_ = 0.040) (Fig. 2B, D; Supplement S9). The response to heat shock across the three regimes followed a similar qualitative pattern across the two sexes, with the S regime showing the strongest reduction in fertility, and the N regime showing weak and non-significant responses. This could imply that sexual selection targeting male postcopulatory reproductive traits in the S regime has led to correlated evolution of female fertility and TSF. Alternatively, it is possible that female TSF evolved independently in the mating regimes yet resulted in a mostly parallel response to that seen in males. Based on the eight line means, we found a marginally non-significant positive genetic correlation between male and female TSF (r = 0.65, p = 0.08, Fig. 4D), which, given the amount of measurement error, does not provide conclusive evidence in favour of one hypothesis over the other.

### Transgenerational effects

Heat shock experienced by fathers, but not mothers, negatively affected offspring fertility (reduction in offspring fertility due to heat shocked father: 7.1 [3.2, 10.5], p_MCMC_ = 0.002; reduction in offspring fertility due to heat shocked mother: -0.5 [-4.19, 2.93], p_MCMC_ = 0.96; interaction parental sex:parental treatment: p_MCMC_ = 0.012, Fig. 5; Supplement S11). We found no evidence that heat shock experienced by either parent would affect offspring TSF, providing no support for adaptive transgenerational plasticity (Supplement S11). We note that the power to detect significant higher-order interactions is modest with these data.

## Discussion

Increased investment into reproduction fuelled by sexual selection is expected to trade-off with fitness components under natural selection (Folstad & Karter, 1992; Zahavi, 1975). However, while postcopulatory sexual selection is widespread in promiscuous species (Parker, 1970), empirical evidence documenting trade-offs between different sperm traits (Pizzari and Parker, 2009; Snook, 2005, Simmons & Fitzpatrick 2012), or between sperm traits and somatic traits (Simmons & Emlen 2006; Lüpold et al., 2015; Parker and Pizzari, 2010; Simmons et al., 2017), is mixed and relatively scarce. One possible explanation could be that optimal allocation strategies often are context-dependent (Harshman & Zera 2007; Ferenci, 2016; Flatt, 2020; Messina and Fry, 2003). Here we first provided simple theoretical arguments to why male adaptation enhancing postcopulatory reproductive success may be associated with a reduction in fertility that manifests at stressful temperatures, and then used long-term experimental evolution in a model species for sexual selection to provide empirical support for this prediction. Specifically, our model and empirical data suggest that reductions in fertility imposed by environmental stress will be most pronounced in highly polyandrous species that have evolved in a constant and benign environment prior to the abrupt environmental change (Figs. 1 & 2, Supplement S1).

What might the long-term consequences of these effects be for polyandrous species facing the increased incidence of heat waves projected under future climate change? Quantitative genetic models of adaptation suggest that sexual selection can improve population viability (Agrawal, 2001; Lorch et al., 2003; Siller, 2001). However, such conclusions rely on two main assumptions.

First, genetic variation for traits under selection needs to be abundant, and population size sufficiently large, to sustain population growth during environmental change (Bürger & Lynch, 1995; Lande & Shannon, 1996). Whether or not fitness-related traits typically harbour sufficient genetic variation to permit the rapid evolution that climate change demands is under debate (Angert et al., 2020; Bonnet et al., 2022; Kokko et al., 2017; Lancaster et al., 2022) and will likely differ between traits and the type of environmental change imposed (Agrawal & Whitlock, 2010; Caruso et al., 2017; null Hoffmann & Merilä, 1999; Rowiński & Rogell, 2017). While male reproductive traits can evolve rapidly (Haerty et al., 2007; Swanson & Vacquier, 2002), recent studies indicate that the evolutionary potential of thermal tolerance is limited (Castañeda et al., 2019; Debes et al., 2021; Kellermann & Heerwaarden, 2019; Morgan et al., 2020; Zwoinska et al., 2020). Moreover, a recent meta-analysis has suggested that the strength of purifying selection increases at elevated temperature, implying that populations facing climate warming will experience an increase in genetic load (Berger et al., 2021). Heat-induced fertility costs associated with postcopulatory sexual selection may therefore have severe repercussions in small populations with limited standing genetic variation in reproductive phenotypes.

Second, quantitative genetic models that predict population-level benefits of sexual selection assume that male adaptation under sexual selection also improves female fitness components (the “genic capture” hypothesis: Rowe and Houle, 1996; Tomkins et al., 2004). However, sexual selection in males can promote genes with detrimental effects on female fitness (Bonduriansky & Chenoweth, 2009). In our experiment, S females, who did not experience fecundity selection themselves, showed high fertility under benign conditions but suffered more from heat stress (Fig 2). While we cannot exclude that some of these effects may have been caused by direct selection on S females via mate choice processes (Hare & Simmons, 2019), our results seem more consistent with sexual selection in males targeting genes that shift allocation away from maintenance towards reproduction in females, with detrimental effects evident under adult heat stress. As female fertility is typically a more important determinate of demography than male fertility (Caswell, 2006; Manning, 1984), this mechanism could contribute further to population decline and extinction threats in warming climates. Nevertheless, experimental studies have illustrated that sexual selection has the potential to aid adaptation to stressful environments in general (reviewed in Cally et al., 2019), and to warm developmental temperatures in particular (Godwin et al., 2020; Parrett & Knell, 2018; Plesnar-Bielak et al., 2012), although the roles of pre-versus postcopulatory sexual selection in driving these patterns remain unclear. Thus, by showing that sperm competition is associated with immediate costs in populations experiencing acute adult heat stress, our results add to a growing body of literature illustrating that sexual selection can impact evolutionary potentials under environmental change (e.g., Fox et al., 2019; García-Roa et al., 2020; Martínez-Ruiz and Knell, 2017; Martinossi-Allibert et al., 2019a; Pilakouta and Ålund, 2021; Rowe and Rundle, 2021; Singh and Agrawal, 2022; Yun et al., 2017). The upcoming challenge is to translate results such as ours into practical insights that will help predict species vulnerability and adaptive potential in the face of climate change.

### Mechanistic explanations

What may be the underlying mechanistic link between sexual selection and TSF? Several transcription factors regulating heat shock protein (hsp) expression have been found to play important regulatory roles during spermatogenesis at benign temperatures (Shiraishi, 2016; Widlak & Vydra, 2017), and male germ cells show a distinct heat stress response compared to other cells (Kim et al., 2013; Michaud et al., 1997; Sarge, 1995). These findings suggest that elements of the heat stress response are employed during spermatogenesis and provide a possible functional basis by which postcopulatory sexual selection may optimise sperm competitive ability at a cost of increased TSF (Dowling & Simmons, 2009). Moreover, reproduction has been shown to generally trade-off with HSP expression (Sørensen et al., 2003), suggesting that this functional basis also may underly the observed increase in TSF of S females (Rodrigues et al., 2022).

For the correlations between male TSF and the previously assayed traits measuring different aspects of sperm competition success, we found no association between TSF and sperm production, suggesting that a simple trade-off between sperm number and quality is unlikely to explain the increased TSF in the S regime. We also found no correlation between TSF and sperm defence (P1; male is first to mate), but a strong a relationship with sperm offence (P2; male is second to mate). Success in sperm defence may to large extent depend on a male’s ability to induce early reproduction in the female (securing paternity of the fraction of eggs fertilized prior to the female mating with another male), while work in fruit flies suggests that success in sperm offence depends on allocation to production of ejaculate components that cause females to dump sperm of the first male from their reproductive tract (Lüpold et al., 2016; McDonough-Goldstein et al., 2022; Wigby et al., 2020). The male ejaculate in *C. maculatus* contains a rich mix of components of which some are thought to be toxic but important in sperm competition (e.g., Yamane et al., 2015), and male genotypes that are successful in sperm competition have been shown to sire offspring of lower quality (Bilde et al., 2009). It is thus possible that the increased TSF in S males could in part have been mediated by the transfer of such toxic components to their female mating partner. Future studies are needed to reveal the exact mechanistic basis behind our results.

### Effects of male rivals

We also investigated plastic responses of TSF to male-male interactions, which typically incurs considerable costs in *C. maculatus* males, as reflected by shortened lifespan (Maklakov & Bonduriansky, 2009). Our data show a tendency for more detrimental effects of male-male interactions in the S and N+S males compared to N males, which could be a sign of adaptation to, and associated costs of, sexual selection. However, while environmental stressors sometimes exacerbate each other’s effects (Relyea & Mills, 2001; Sejian et al., 2011), we found no such obvious effects here for heat stress and male competition. The presence of male rivals do not always favour increased allocation to sperm production but can in certain scenarios, where such interactions confer considerable costs, instead favour reduced reproductive effort in favour of maintenance (Parker, 1990; Parker & Pizzari, 2010). It might thus be that several simultaneous effects triggered by male rivals (i.e., overall reduction in condition coupled with shifts in germline allocation) could have counteracting effects on male TSF. Additionally, effects of male-male interactions on TSF were only monitored in the first mating, where overall effects of heat shock were weak, limiting our inferences.

### Transgenerational effects

The full consequence of heat waves on population viability will depend on if and how effects in exposed parents get transferred to offspring, where detrimental effects at the population-level can be exacerbated via further reductions in the quality and fertility of their surviving offspring. Exposed parents may also prime their gametes with epigenetic information helping offspring to better cope with future heat stress. Indeed, such adaptive trans-generational plasticity can provide an avenue to maintain population fitness under climate change (Bonduriansky & Day, 2009; Donelson et al., 2018; A. A. Hoffmann & Sgrò, 2011). We found that in *C. maculatus*, paternal heat shock reduces offspring fertility while mothers transferred no obvious effects to offspring. This corroborates findings of a recent study by Sales et al. (2018), in which heat exposure of paternal sperm (either via the father himself or via the inseminated mother) resulted in decreased survival and fitness of offspring. We found no indication that offspring of heat-exposed fathers performed better under heat stress conditions relative to controls from untreated parents, suggesting that adaptive transgenerational plasticity is unlikely to remedy fertility loss due to heat stress. Strikingly, in our transgenerational experiment there was no apparent fertility reduction detected at all in the exposed F0 fathers (Fig. 3; first mating for isolated N+S males) while the offspring deriving from this mating suffered a 10% reduction in fertility on average (Fig. 5, bottom panels). This result is similar to that reported recently for field crickets (Simmons et al., 2022) and highlights that heat waves can have long-lasting effects in natural populations that may remain undetected in experimental studies unless appropriate designs are used.

## Conclusions

Here we have provided evidence for fertility trade-offs associated with adaptation under post-copulatory sexual selection. Our empirical data and simple model suggest that such trade-offs may become more apparent under environmental stress because strong directional selection for male traits that increase postcopulatory reproductive success in benign conditions may lead to allocation strategies that have detrimental effects on fertility once environmental conditions worsen and put larger demands on germline maintenance and repair. This fertility debt owing to sexual selection may have particularly detrimental effects in the light of the findings that sexual selection also affected female TSF, and that effects of paternal heat shock permeate through generations in *C. maculatus*. The increase in heat waves expected under climate warming may thus cause pronounced reductions in population size in species evolving under postcopulatory sexual selection that may elevate their extinction risk unless standing genetic variation for heat tolerance is abundant.

## Acknowledgements

This research was funded by grants from the Swedish research council (VR: 2019-05024) and Carl Tryggers Stiftelse (CTS:18:32) to D.B..

## Author contributions

J.B., M.Z., M.K., and D.B. conceived and designed the study. J.B., M.Z., and M.K. performed the experiments. J.B., M.Z., and D.B. analysed the data, and J.B. and D.B. wrote the manuscript. All authors commented on manuscript drafts.

## Data Accessibility

All data generated during the experiments presented in this study will be made publicly available upon acceptance.

