## Supplementary material for "Heat stress reveals a fertility debt owing to postcopulatory sexual selection"

**Supplementary note S1:**

*A heuristic model of postcopulatory sexual selection and environmental sensitivity of fertility*

We modelled the environmental sensitivity of fertility arising from reproduction-maintenance trade-offs in the germline by employing a life-history framework and a version of the “Y-model” of resource acquisition and allocation^1,2^. The aim was to provide general and qualitative predictions for how a long history of strong postcopulatory sexual selection would affect male fertility under abrupt environmental change. We therefore constructed a very simple model with as few assumptions as possible. We did not consider trade-offs with somatic performance or trade-offs between current and future reproduction. Moreover, as the focus was on predicting how evolutionary history would affect immediate responses to future climate change (e.g. heat waves), we did not model adaptive plasticity in germline allocation (optimal strategies were assumed to be fixed), or further evolution following the environmental change.

Individual condition (*C*) was assumed to determine the amount of resources that can be allocated to germline maintenance (*M*) in form of anti-oxidative defence and repair needed to maintain ejaculate quality and gamete viability^3,4^, or reproductive effort (*R*) in form of production of gametes and ejaculatory components that increase a male’s success in sperm competition, such that:

*C = R + M;	R = kC;	M =* (1-*k*)*C*.	**Eq. 1**

where *k* is the proportion of resources allocated to reproduction. Gamete viability,$\zeta$, is assumed to be dependent on the amount of maintenance per reproductive effort (i.e. per gamete or ejaculate volume):

$\zeta\approx\left[ \frac{\left( 1-k \right)}{\left( 1+k \right)} \right]^{a}$	**Eq. 2**

where $a$ describes environmentally dependent consequences of sub-maximal germline maintenance, such that some environmental conditions will impair fertility more than others (e.g. hot temperature or high salinity increases $a$). The addition of +1 to the denominator of Eq. 2 assures that viability ranges between 0 and 1, but we note that this choice was arbitrary and that other expressions for Eq. 2 resulted in the same qualitative results. If reproductive success follows a power function of reproductive effort, and if fitness, $\omega$, is the product of sperm competition success and gamete viability, then:

$\omega\approx R^{b}{\zeta=(kC)}^{b}\left[ \frac{\left( 1-k \right)}{\left( 1+k \right)} \right]^{a}$	**Eq.3**

where parameter $b$ describes how reproductive effort translates into postcopulatory reproductive success. When $b$ < 1, reproductive success is less than proportional to investment and postcopulatory sexual selection is relatively weak, whereas $b$ > 1 gives a disproportionate advantage to individuals investing more in reproductive effort and sexual selection is strong. This also means that individuals in a population experiencing strong sexual selection need to invest much more in reproductive effort to secure a significant share of paternity relative to individuals from a population where sexual selection is weak. Given a trade-off between investment in reproductive effort and germline maintenance (Eq. 1), such excessive germline allocation is predicted to result in reduced fertility, an effect that is particularly pronounced under harsh environmental conditions (Fig. S1.1).

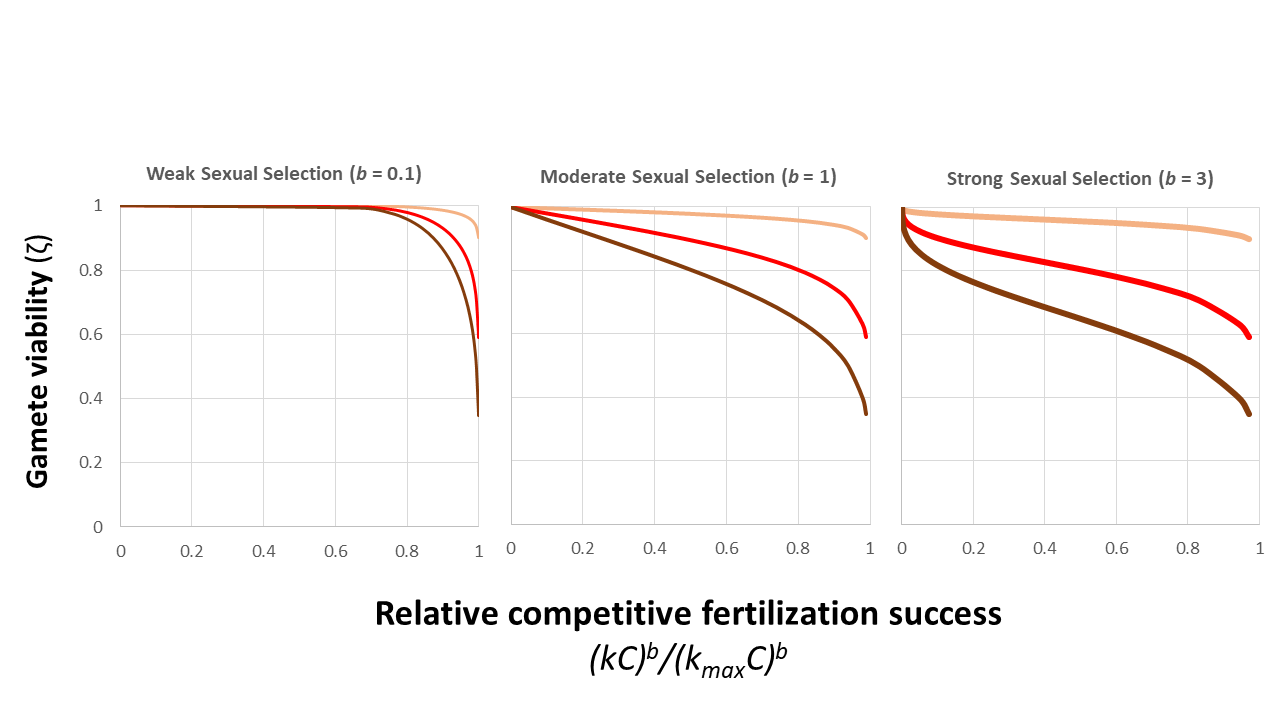

**C**

**B**

**A**

**Supplementary Figure S1.1.** The relationship between competitive reproductive success (i.e. success in sperm competition) and gamete viability for different strengths of sexual selection (*b* = 0.1; 1; 3 from panel **A-C**) and viability selection (*a*; salmon = 0.02, red = 0.10, brown = 0.20). Competitive reproductive success is expressed relative to the success of an individual with the same condition (*C*) investing all resources in reproduction (*k_max_* = 1). Individuals experiencing strong sexual selection (panel **C**) need to invest more in reproduction to get the same share of paternity compared to individuals experiencing weak sexual selection (panel **A**) and pay a fertility cost in terms of reduced gamete viability. This cost becomes more pronounced in harsh environments (brown lines).

The optimal germline allocation strategy (*k*_opt_) for different strengths of sexual selection (*b*) and viability selection (*a*) is given by differentiation of equation (3) with respect to *k*:

$k_{opt}=\frac{\sqrt{b^{2}+a^{2}}-a}{b}$	**Eq. 4**

which shows that optimal allocation to reproduction, $k_{opt}$, increases with the strength of sexual selection, $b$, and decreases with environmentally dependent viability selection, $a$. Unsurprisingly, increased sexual selection does indeed lead to decreased gamete viability as follows from the trade-off scenario described by equation (1) (Supplementary Fig. S1.1 and Fig. S1.2A). Optimal allocation (and resulting fertility) is independent of condition, $C$, when reproductive success is a power function of investment, but we note that reproductive effort and gamete viability can either be increasing or decreasing functions of condition, depending on the fitness functions used (results not shown, but see: ^5–7^).

What consequences do differences in mating system and a history of intense sperm competition (high $b$) have for fertility responses to increased environmental stress (increases in $a$)? We illustrate these effects by first replacing $a$ in equation (4) with $a_{anc}$, representing viability selection in a relatively benign ancestral environment, making it possible to solve for $k_{opt}$ for different values of $b$. We then replace *k* in equation (2) with this expression for $k_{opt}$ and differentiate with respect to $a$ to show how gamete viability, $\zeta$, is affected by increasing environmental stress for allocation strategies that have evolved under different scenarios of sexual selection and viability selection in the ancestral environment:

$\frac{d\zeta}{da}= \left[ \frac{\sqrt{b^{2}+a_{anc}^{2}}-b}{a_{anc}} \right]^{a}.\ln\left[ \frac{\sqrt{b^{2}+a_{anc}^{2}}-b}{a_{anc}} \right]$	**Eq. 5**

Predictions from equation (5) are presented in Supplementary Fig. S1.2B (see also Fig. 1B in main text) and show that, for any strength of gamete viability selection in the ancestral environment, populations that have evolved under a history of strong sexual selection are predicted to suffer a greater fertility loss following increased environmental stress.

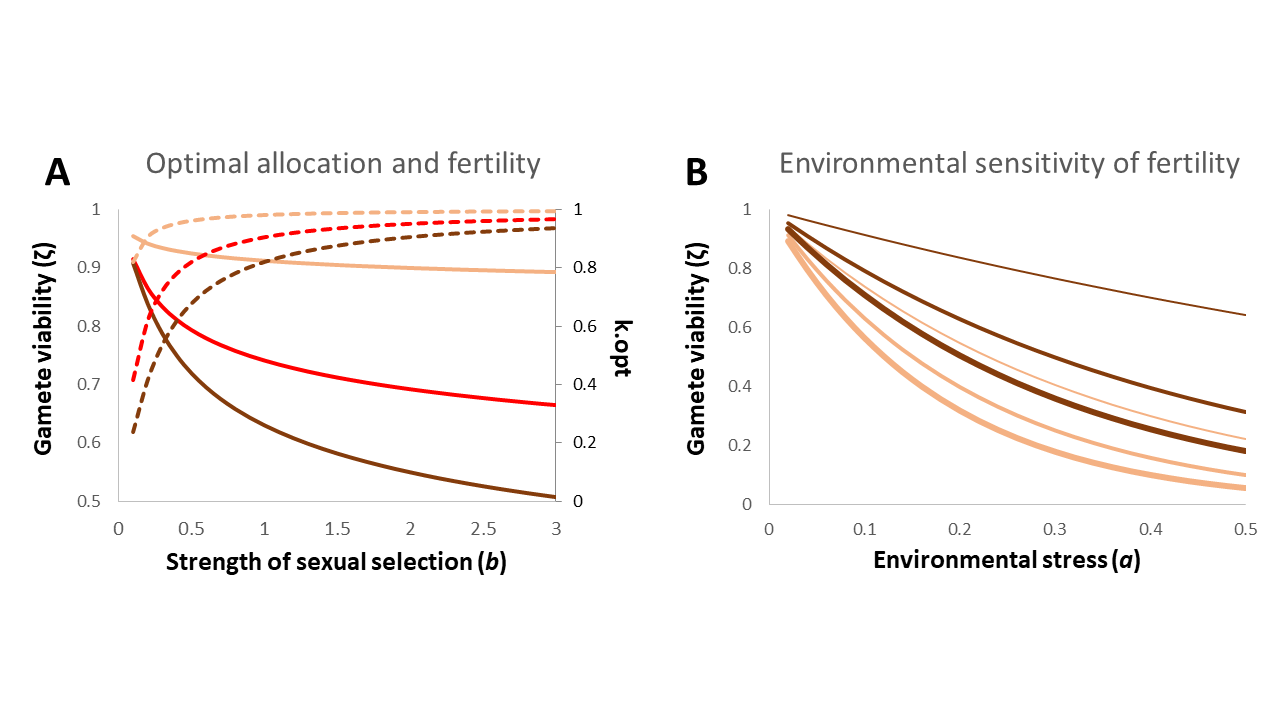

**Supplementary Figure S1.2.** In **A**) the optimal reproductive effort ($k_{opt}$; hatched lines) and resulting gamete viability ($\zeta$; full lines) for different levels of sexual selection (*b*) and viability selection (*a*; salmon = 0.02, red = 0.10, brown = 0.20). In **B**) the change in gamete viability as environmental stress (*a*) changes from ancestral conditions ($a_{anc}$; salmon = 0.02, brown = 0.20) for populations that have evolved optimal allocation under either weak (*b* = 0.1, thin lines), moderate (*b* = 1, intermediate lines) or strong (*b* = 3, thick lines) postcopulatory sexual selection.

We note that these are qualitative predictions and that the use of different fitness- and trade-off functions change results quantitatively. We also note that, because (postcopulatory) sexual selection is inherently a frequency dependent process^8^, it is likely that abrupt changes in the environment (increases in *a*) could themselves modulate parameter *b*, by for example changing the density and quality of rivalling males at mating sites^5,8–11^. Such dynamics would ultimately need to be considered in more sophisticated models concerned with how germline plasticity and future evolution of reproductive strategies in changing environments would affect fertility and population health. Here, however, we kept *b* constant to describe the strength of sexual selection in the ancestral environment as we were interested in generating predictions for immediate fertility responses under abrupt environmental change attributed to the organism’s evolutionary history of natural and sexual selection. We then could compare these qualitative predictions directly with our empirical data (see main text).

**Supplementary figure S2:**

*Experimental design: main experiment*

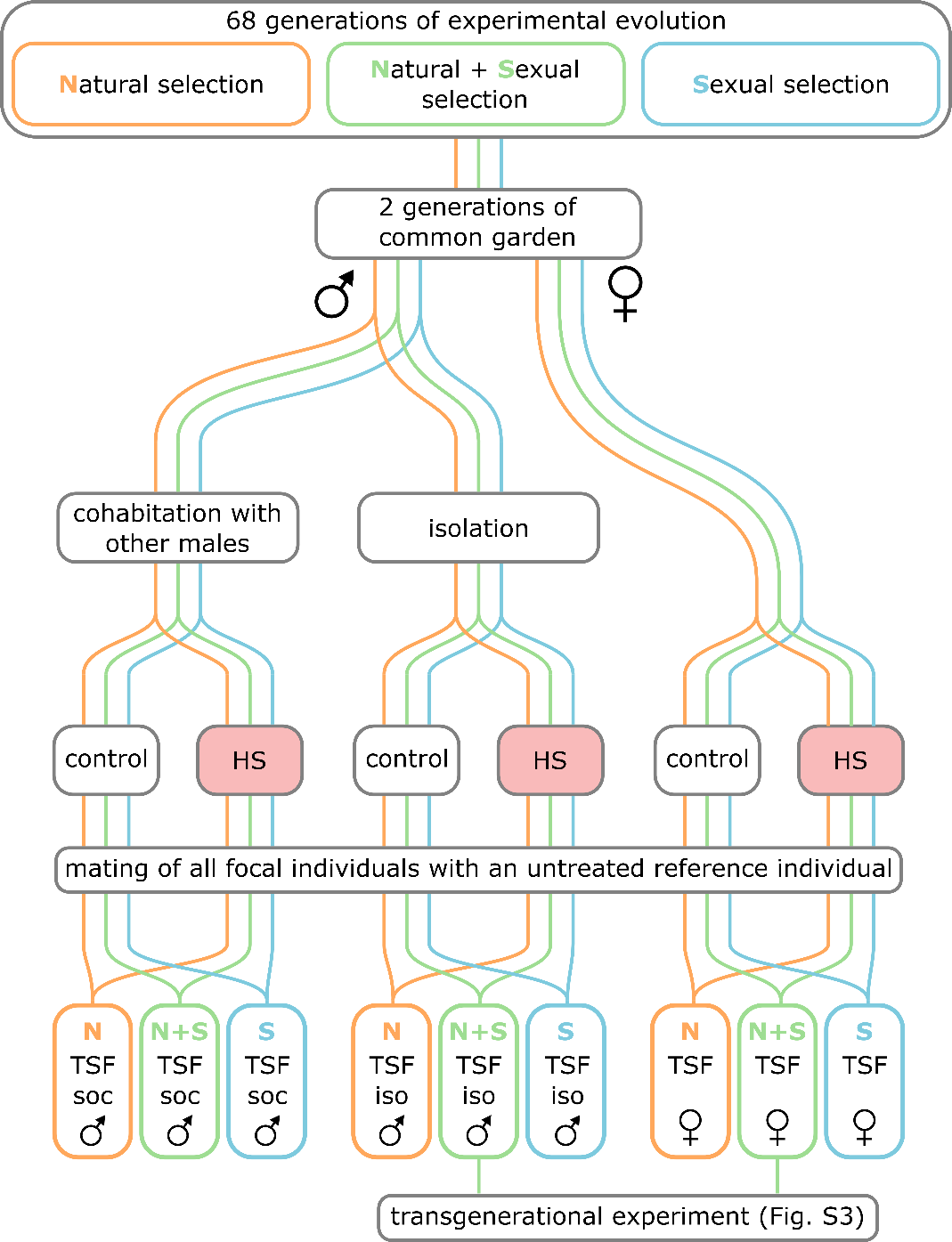

**Design of the main experiment, including experimental evolution under the three alternative selection regimes, and two generations of common garden prior to the experimental generation.** In the first experimental generation (parental), virgin beetles of all three regimes were picked within 24 hours after eclosion and males were either kept isolated or kept together with other males. Subsequently, half of the beetles of each regime and treatment combination were exposed to a heat shock. The thermal sensitivity of fertility was assessed by comparing heat shocked and control beetles within regime and treatment according to: TSF = 1 – (offspring_heat shocked_ / offspring_control_). F1 offspring of heat shocked and control N+S males and females were used for the assessment of transgenerational effects of heat shock (see supplementary figure S4).

**Supplementary figure S3:**

*Daily temperature curve for* Lomé*, Togo*

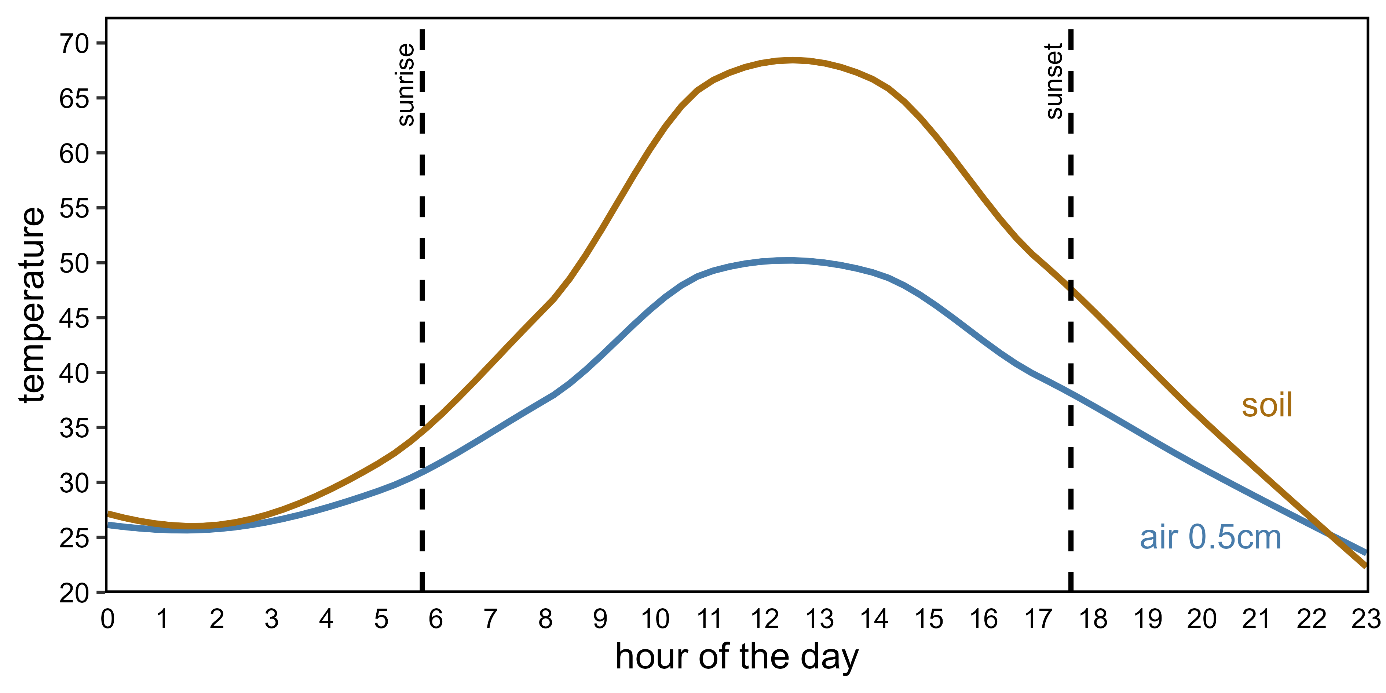

**Soil and air temperature curve as it may occur on a day in November, the hottest month of the year, in Lomé, Togo (06°10#N 01°13#E), the original collection site of the beetle stock used here.** The presented temperatures have been estimated using NicheMapR (version 3.2.0; Kearney and Porter, 2016, Ecography), an R software package interpolating climate data in order to model microclimates on a very fine scale. The shown curves represent soil temperature and air temperature in a height of 0.5cm. We assumed a climate change scenario of global warming of 1.5°C, a day with no cloud coverage, even ground, average local wind speed, 15% shaded area, and conservative values for soil albedo (15%) and emissivity (0.9).

**Supplementary table S4:**

*Sample sizes*

Table S2.1: Sample size per cell for analysis of male thermal sensitivity of fertility

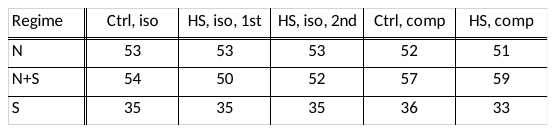

Table S2.2: Sample sizes per cell for analysis of female thermal sensitivity of fertility

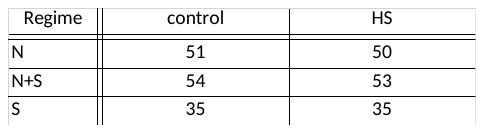

Table S2.3: Sample sizes per cell for analysis of transgenerational effects

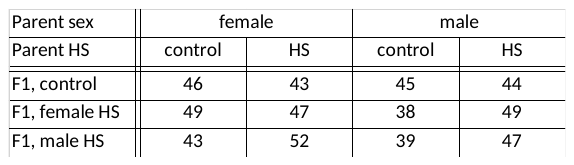

**Supplementary material S5:**

*Effect of multiple mating in the S regime.*

The strongest decline in fertility was observed following the second mating of heat shocked S males. We here show that S males do not show such a decline in fertility if kept under benign conditions, and that this effect can be assigned to the heat shock treatment. We ran a follow-up experiment in which we allowed males (N=49) from the S regime kept at benign conditions to mate two times with 6 hours in between matings, as was done for males that had been heat shocked in the main experiment.

We first analysed differences in fertility between first and second matings of S males kept under benign conditions (i.e., the data collected during the follow-up experiment). We then also included data from the main experiment on fertility of isolated control males (first mating) and from the second mating of isolated heat shocked males. The term [experiment] represents from which experiment the males were taken and the interaction term [mating:experiment] tests the statistical significance of heat shock in the second mating.

*Effects of mating number in males kept in benign conditions (Follow-up):*

glm(offspring~ mating*line , family ="quasipoisson", data = followUp[follow$experiment == “followUp”,])

| Analysis of Deviance Table (Type III tests) | | | |
| --- | --- | --- | --- |
|  | Χ^2^ | df | p-value |
| mating | 2.41 | 1 | 0.122 |
| line | 0.08 | 1 | 0.77 |
| mating:line | 0.73 | 1 | 0.3917 |

*Effects of mating number in males exposed to heat shock (main experiment):*

glm(offspring~ mating*line , family ="quasipoisson", data = followUp[follow$experiment == “Main”,])

| Analysis of Deviance Table (Type III tests) | | | |
| --- | --- | --- | --- |
|  | Χ^2^ | df | p-value |
| mating | 9.16 | 1 | **0.002**** |
| line | 0.02 | 1 | 0.88 |
| mating:line | 0.24 | 1 | 0.62 |

*Effects of heat shock on fertility decline:*

glm(offspring~ mating*experiment*line , family ="quasipoisson", data = follow2)

| Analysis of Deviance Table (Type III tests) | | | |
| --- | --- | --- | --- |
|  | Χ^2^ | df | p-value |
| mating | 2.47 | 1 | 0.12 |
| experiment | 1.10 | 1 | 0.29 |
| line | 0.12 | 1 | 0.73 |
| mating:experiment | 5.77 | 1 | **0.016*** |
| mating:line | 0.91 | 1 | 0.34 |
| experiment:line | 0.02 | 1 | 0.88 |

**Supplementary figure S6:**

*Experimental design: transgenerational experiment*

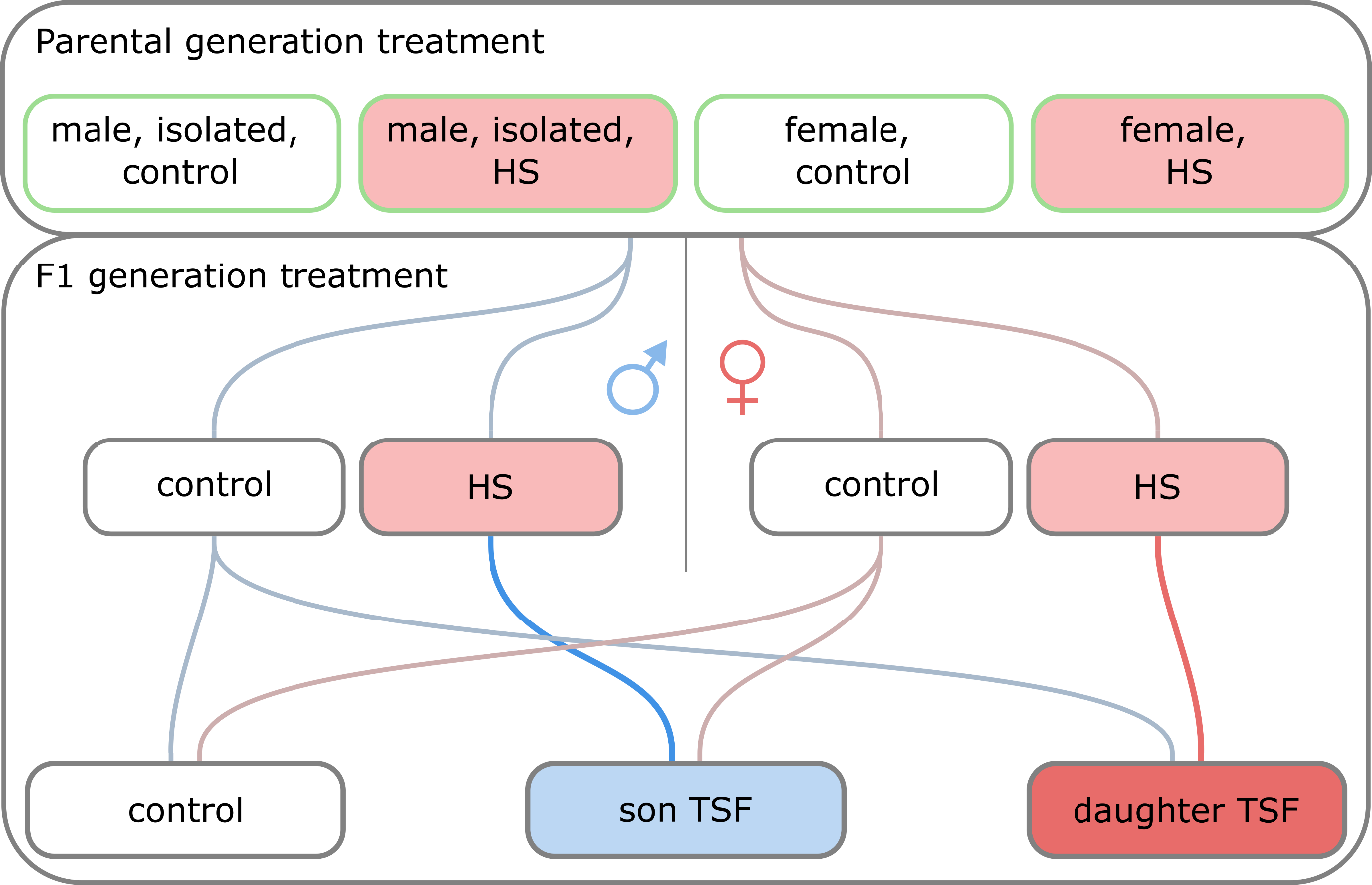

**The experimental design used to assess transgenerational effects in offspring of heat shock in male and female parents.** Focal parents were from N+S lines and were either control individuals or had been exposed to heat shock in the first generation of the experiment. Maternal and paternal transgenerational effects of the parental heat shock treatment was assessed by measuring the fertility of F1 offspring mating pairs. To assess whether the parental heat shock influenced the TSF of F1 sons and daughters, we also applied heat shock to the F1 generation in a sex-specific manner before assaying fertility. All offspring crosses were performed within line replicate and parental treatment group, while avoiding sib-mating. We used the same experimental procedure as in the main experiment for the heat treatment and fitness assays.

**Supplementary table S7:**

*Full model specifications and output of the MCMCglmm used to analyse thermal sensitivity of fertility of the first and second mating of males kept in isolation prior to the heat shock application.*

For this analysis heat shock treatment and mating number were combined in one factor with three levels (control, heat shock first mating (1h after heat shock), and heat shock second mating (7h after heat shock)).

Data: Male TSF data, excluding males that were exposed to socio-sexual competition (see S2.1, excluding “Ctrl, comp” and “HS, comp”).

prior_male = list(R = list(V = diag(9), nu = 10ˆ-6), G = list(G1 = list(V = 1, nu = 10ˆ-6), G2 = list(V = 1, nu = 10ˆ-6), G3 = list(V = 1, nu = 10ˆ-6), G4 = list(V = 1, nu = 10ˆ-6), G5 = list(V = 1, nu = 10ˆ-6)))

MCMCglmm(offspring ~ regime*treatment, random = ~line + treatment:line + block + treatment:block + ID, rcov = ~ idh(treatment:regime):units, data = F1males[F1males$competition == "No",], family = "poisson", prior = prior_male, nitt=2200000, slice=TRUE, burnin=200000, thin=2000, verbose = FALSE)

| Fixed effects: |  |  |  |  |
| --- | --- | --- | --- | --- |
|  | posterior mean | lower 95% CI | upper 95% CI | p_MCMC_ |
| regimeN - Control (Intercept) | 4.169 | 4.051 | 4.293 | **<0.001***** |
| regimeS | 0.063 | -0.063 | 0.173 | 0.244 |
| regimeNS | 0.084 | -0.044 | 0.198 | 0.178 |
| treatmentHS1 | -0.134 | -0.348 | 0.068 | 0.230 |
| treatmentHS2 | -0.064 | -0.204 | 0.075 | 0.346 |
| regimeS:treatmentHS1 | -0.069 | -0.343 | 0.207 | 0.618 |
| regimeNS:treatmentHS1 | 0.146 | -0.064 | 0.371 | 0.198 |
| regimeS:treatmentHS2 | -0.260 | -0.489 | -0.037 | **0.014*** |
| regimeNS:treatmentHS2 | -0.01 | -0.152 | 0.102 | 0.756 |

| Random effects: |  |  |  |
| --- | --- | --- | --- |
|  | posterior mean | lower 95% CI | upper 95% CI |
| line | 0.0016 | 2.69E-07 | 0.0070 |
| treatment:line | 0.0002 | 1.12E-07 | 0.0010 |
| block | 0.0028 | 2.08E-07 | 0.0120 |
| treatment:block | 0.0075 | 7.14E-07 | 0.0184 |
| ID | 0.0064 | 1.10E-07 | 0.0174 |

**Supplementary table S8:**

*Full model specifications and output of the MCMCglmm used to analyse effects of male-male competition on thermal sensitivity of fertility of the first ejaculate.*

Data: Male TSF data, only including data on first ejaculates of isolated males (see S2.1, excluding “HS, iso, 2^nd^”).

Prior_comp = list(R = list(V = diag(12), nu = 10ˆ-6), G = list(G1 = list(V = 1, nu = 10ˆ-6), G2 = list(V = 1, nu = 10ˆ-6), G3 = list(V = 1, nu = 10ˆ-6), G4 = list(V = 1, nu = 10ˆ-6), G5 = list(V = 1, nu = 10ˆ-6), G5 = list(V = 1, nu = 10ˆ-6)))

MCMCglmm(offspring ~ regime*HS*competition, random = ~line + HS:line + block + HS:line:competition + HS:block + line:competition, rcov = ~ idh(HS:competition:regime):units, data = F1males[F1males$treatment != "IsoHS2",], family = "poisson", prior = prior_comp, nitt=2200000, slice=TRUE, burnin=200000, thin=2000, verbose = FALSE)

Excluding males that failed to mate:

| Fixed effects: |  |  |  |  |
| --- | --- | --- | --- | --- |
|  | posterior mean | lower 95% CI | upper 95% CI | p_MCMC_ |
| regimeN (Intercept) | 4.167 | 4.045 | 4.295 | **<0.001***** |
| regimeS | 0.069 | -0.076 | 0.206 | 0.308 |
| regimeNS | 0.086 | -0.044 | 0.241 | 0.194 |
| HSYes | -0.130 | -0.338 | 0.061 | 0.207 |
| competitionYes | -0.019 | -0.125 | 0.087 | 0.692 |
| regimeS:HSYes | -0.080 | -0.366 | 0.201 | 0.569 |
| regimeNS:HSYes | 0.142 | -0.091 | 0.349 | 0.204 |
| regimeS:competitionYes | -0.076 | -0.208 | 0.062 | 0.231 |
| regimeNS:competitionYes | -0.170 | -0.360 | 0.031 | **0.083** |
| HSYes:socioYes | 0.111 | -0.097 | 0.301 | 0.261 |
| regimeS:HSYes:competitionYes | 0.097 | -0.222 | 0.372 | 0.511 |
| regimeNS:HSYes:competitionYes | -0.020 | -0.297 | 0.231 | 0.895 |

| Random effects: |  |  |  |
| --- | --- | --- | --- |
|  | posterior mean | lower 95% CI | upper 95% CI |
| line | 0.0015 | 1.49E-07 | 0.0066 |
| HS:line | 0.0011 | 1.13E-07 | 0.0052 |
| block | 0.0031 | 1.46E-07 | 0.0115 |
| HS:line:competition | 0.0002 | 8.17E-08 | 0.0014 |
| HS:block | 0.0045 | 1.43E-07 | 0.0116 |
| line:socio | 0.0004 | 1.01E-07 | 0.0018 |

Including males that failed to mate as couples with offspring equal to zero:

| Fixed effects: |  |  |  |  |
| --- | --- | --- | --- | --- |
|  | posterior mean | lower 95% CI | upper 95% CI | p_MCMC_ |
| regimeN (Intercept) | 4.175 | 3.984 | 4.368 | **<0.001***** |
| regimeS | 0.065 | -0.067 | 0.182 | 0.270 |
| regimeNS | 0.088 | -0.035 | 0.226 | 0.172 |
| HSYes | -0.203 | -0.524 | 0.103 | 0.187 |
| socioYes | -0.017 | -0.131 | 0.077 | 0.720 |
| regimeS:HSYes | -0.001 | -0.309 | 0.285 | 0.983 |
| regimeNS:HSYes | 0.116 | -0.186 | 0.353 | 0.4393 |
| regimeS:socioYes | -0.078 | -0.198 | 0.054 | 0.244 |
| regimeNS:socioYes | -0.180 | -0.385 | 0.001 | **0.058** |
| HSYes:socioYes | 0.009 | -0.284 | 0.303 | 0.934 |
| regimeS:HSYes:socioYes | 0.058 | -0.297 | 0.454 | 0.761 |
| regimeNS:HSYes:socioYes | 0.133 | -0.219 | 0.519 | 0.488 |

| Random effects: |  |  |  |
| --- | --- | --- | --- |
|  | posterior mean | lower 95% CI | upper 95% CI |
| line | 0.0006 | 1.19E-07 | 0.0029 |
| HS:line | 0.0012 | 1.80E-07 | 0.0061 |
| block | 0.0059 | 1.23E-07 | 0.0287 |
| HS:line:socio | 0.0003 | 1.97E-07 | 0.0017 |
| HS:block | 0.0346 | 0.005 | 0.0743 |
| line:socio | 0.0003 | 9.11E-08 | 0.0012 |

**Supplementary table S9:**

*Full model specifications and output of the MCMCglmm used to analyse thermal sensitivity of fertility of females.*

Data: Female TSF data (see S2.2, all data).

prior_female = list(R = list(V = diag(6), nu = 10ˆ-6), G = list(G1 = list(V = 1, nu = 10ˆ-6), G2 = list(V = 1, nu = 10ˆ-6), G3 = list(V = 1, nu = 10ˆ-6), G4 = list(V = 1, nu = 10ˆ-6)))

MCMCglmm(offspring ~ regime*HS, random = ~line + HS:line + block + HS:block, rcov = ~ idh(regime:HS):units, data = females, family = "poisson", prior = prior_female, nitt=2200000, slice=TRUE, burnin=200000, thin=2000, verbose = FALSE, pr = TRUE)

| Fixed effects: |  |  |  |  |
| --- | --- | --- | --- | --- |
|  | posterior mean | lower 95% CI | upper 95% CI | p_MCMC_ |
| regimeN (Intercept) | 4.0540 | 3.735 | 4.345 | **<0.001***** |
| regimeS | 0.366 | 0.044 | 0.721 | **0.032*** |
| regimeNS | 0.337 | -0.0061 | 0.666 | **0.0456** |
| HSYes | 0.226 | -0.156 | 0.517 | 0.196 |
| regimeS:HSYes | -0.477 | -0.907 | -0.0001 | **0.040*** |
| regimeNS:HSYes | -0.300 | -0.677 | 0.097 | 0.142 |

| Random effects: |  |  |  |
| --- | --- | --- | --- |
|  | posterior mean | lower 95% CI | upper 95% CI |
| line | 0.0051 | 1.22e-07 | 0.0237 |
| HS:line | 0.0023 | 1.84e-07 | 0.0129 |
| block | 0.0012 | 1.74e-07 | 0.0066 |
| HS:block | 0.0011 | 2.41e-07 | 0.0059 |

**Supplementary figure S10:**

*Figure of effects of socio-sexual interactions and heat shock equivalent to figure 3 in the main text but including males that failed to mate as zero offspring.*

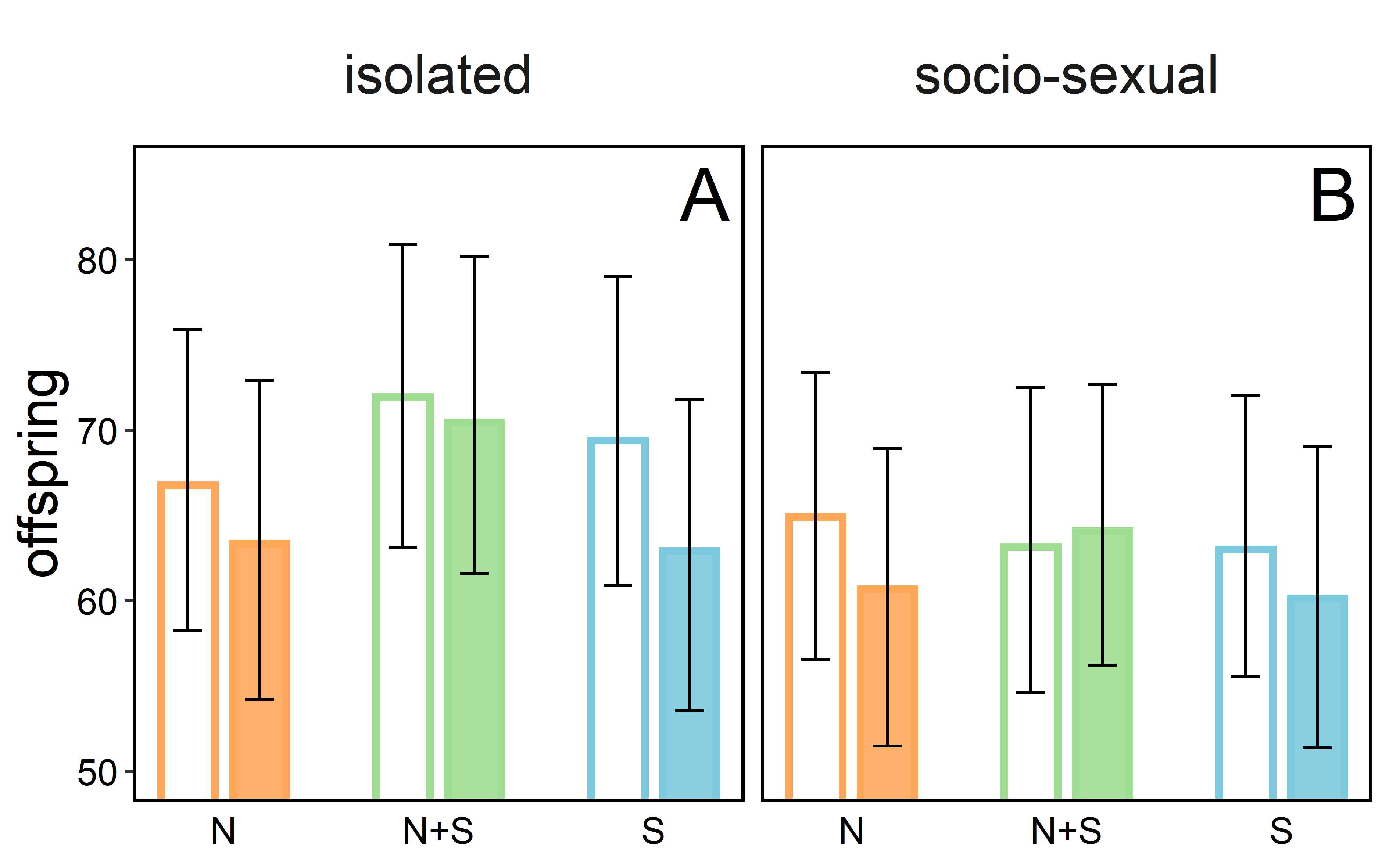

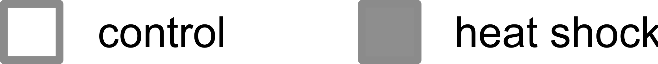

**The effect of male-male interactions on male fertility**. Fertility of couples from the N (orange), N+S (green) and S (blue) regime. Focal males were either kept at benign conditions (open bars) or exposed to heat shock (closed bars) and were either kept isolated or in groups of three prior to heat shock and mating. Bars represent posterior means and whiskers 95% credible intervals.

**Supplementary table S11:**

*Full model specifications and output of the MCMCglmm used to analyse transgenerational effects of a heat shock on offspring quality and thermal sensitivity of fertility.*

This analysis includes the sex of the focal parent (originating from the selection line, while the mating partner originated from the ancestral population) (male/female), parental treatment (control/heat shock), and the treatment of the offspring couple (male heat shock/female heat shock/control).

Data: F2 offspring data (see S2.3, all data).

prior_transgen = list(R = list(V = diag(3), nu = 10^-6), G = list(G1 = list(V = 1, nu = 10^-6),

G2 = list(V = 1, nu = 10^-6), G3 = list(V = 1, nu = 10^-6), G4 = list(V = 1, nu = 10^-6)))

MCMCglmm(offspring ~ P.sex*P.treatment*F1.treatment + line, random = ~ block + P.sex:block + P.treatment:block + P.sex:P.treatment:block, rcov = ~ idh(F1.treatment):units, data = F2data, family = "poisson", prior = priorF2.1, nitt=2200000, slice=TRUE, burnin=200000, thin=2000, verbose = FALSE)

| Fixed effects: |  |  |  |  |
| --- | --- | --- | --- | --- |
|  | posterior mean | lower 95% CI | upper 95% CI | p_MCMC_ |
| P.sexF (Intercept) | 4.280 | 4.208 | 4.346 | **<0.001***** |
| P.sexM | 0.104 | 0.011 | 0.191 | **0.026*** |
| P.treatmentHS | 0.016 | -0.057 | 0.086 | 0.652 |
| F1.treatmentF | -0.008 | -0.082 | 0.064 | 0.804 |
| F1.treatmentM | -0.056 | -0.144 | 0.037 | 0.238 |
| P.sexM:P.treatmentHS | -0.123 | -0.220 | -0.024 | **0.016*** |
| P.sexM:F1.treatmentF | -0.039 | -0.144 | 0.072 | 0.462 |
| P.sexM:F1.treatmentM | 0.029 | -0.114 | 0.151 | 0.640 |
| P.treatmentHS:F1.treatmentF | -0.026 | -0.129 | 0.080 | 0.604 |
| P.treatmentHS:F1.treatmentM | -0.027 | -0.145 | 0.098 | 0.688 |
| P.sexM:P.treatmentHS:F1.treatmentF | 0.081 | -0.064 | 0.233 | 0.302 |
| P.sexM:P.treatmentHS:F1.treatmentM | -0.016 | -0.184 | 0.171 | 0.832 |
| Line 2 | 0.092 | 0.054 | 0.134 | **<0.001***** |
| Line 3 | 0.001 | -0.044 | 0.042 | 0.932 |

| Random effects: |  |  |  |
| --- | --- | --- | --- |
|  | posterior mean | lower 95% CI | upper 95% CI |
| line | 0.0252 | 1.44E-07 | 0.0286 |
| F1.sex:line | 0.0014 | 1.11E-07 | 0.0042 |
| block | 0.0006 | 1.44E-07 | 0.0028 |
| F1.sex:block | 0.0002 | 1.47E-07 | 0.0008 |
| P.sex:P.treatment:F1.sex:line | 0.0004 | 1.01E-07 | 0.0018 |
| P.sex:P.treatment:line | 0.0002 | 1.29E-07 | 0.0010 |
| line:P.sex | 0.0006 | 1.67E-07 | 0.0031 |
